# Isoform-level Ribosome Occupancy Estimation Guided by Transcript Abundance with Ribomap

**DOI:** 10.1101/017509

**Authors:** Hao Wang, Joel McManus, Carl Kingsford

## Abstract

Ribosome profiling is a recently developed high-throughput sequencing technique that captures approximately 30 bp long ribosome-protected mRNA fragments during translation. Because of alternative splicing and repetitive sequences, a ribosome-protected read may map to many places in the transcriptome, leading to discarded or arbitrary mappings when standard approaches are used. We present a technique and software that addresses this problem by assigning reads to potential origins proportional to estimated transcript abundance. This yields a more accurate estimate of ribosome profiles compared with a naϊve mapping. Ribomap is available as open source at http://www.cs.cmu.edu/∼ckingsf/software/ribomap.

## 1. Introduction

Ribosome profiling (ribo-seq) provides snapshots of the positions of translating ribosomes by sequencing ribosome-protected fragments [14]. The distribution of ribo-seq footprints along a transcript, called the ribosome profile, can be used to analyze translational regulation and discover alternative initiation [9], alternative translation and frameshifting [21], and may eventually lead to a better understanding of the regulation of cell growth, the progression of aging [18] and the development of diseases [12, 35]. Different environmental conditions such as stress or starvation alter the ribosome profile patterns [10, 14], indicating possible changes in translational regulation.

In higher eukaryotes, alternative transcription initiation, pre-mRNA splicing, and 3’ end formation result in the production of multiple isoforms for most genes. The resulting isoforms can have dramatically different effects on mRNA stability [19] and translation regulation [33]. However, to date ribosome profiling analyses have been conducted at the gene, rather than isoform, level using either a single ‘representative’ isoform (e.g. [11]) or exon union profiles (e.g. [25]). The lack of isoform-level analysis of ribo-seq data is partially due to the absence of the necessary bioinformatic tools. Here, we present a conceptual framework and software (Ribomap) to quantify isoform-level ribosome profiles. By accounting for multi-mapping sequence reads using RNA-seq estimates of isoform abundance, Ribomap produces accurate isoform-specific ribosome profiles.

The challenge in estimating isoform ribosome profiles is that a short ribo-seq read may map to many different transcripts. Ambiguous mappings are not rare in ribo-seq data and can be caused by either repetitive sequences along the genome or alternative splicing [13]. For example, in the human Hela cell ribo-seq data (GSM546920, [11]), among all mapped reads (about 50% of all reads), only 14*%* can be uniquely mapped to a single location of a single mRNA isoform, 22*%* can be mapped to multiple regions on the reference genome due to repetitive sequences, and 64% can be mapped to multiple mRNAs due to alternative splicing. Ribomap deals with both types of ambiguous mappings, and therefore does not discard multi-mapped reads, resulting in more of the data being used. In this example, the mapping rate of Ribomap is 50*%* compared to 7*%* if only uniquely mapped reads are used.

Estimation of mRNA isoform abundance from RNA-seq has also had to deal with ambiguous mappings [16, 23, 26]. However, unlike in RNA-seq, coverage in ribo-seq is highly non-uniform regardless of sequencing bias since ribosomes move along mRNAs at non-uniform rates, and it is in fact the non-uniformities that are of interest. Further, ambiguous mappings are much worse for ribo-seq data since the read length cannot exceed the ribosome size (approximately 30bp), while paired-end and longer reads can be generated from RNA-seq experiments to reduce the problem of ambiguous mappings. Methods developed for transcript abundance are therefore not applicable to assigning ribo-seq reads.

By observing that ambiguous mappings are mainly caused by multiple isoforms (Supplementary Figure 2), Ribomap assigns ribo-seq reads to locations using estimated transcript abundance of the candidate locations. On simulated data, our approach yields a more precise estimation of ribosome profiles compared with a pure mapping-based approach. Further, the ribosome abundance derived using our method correlates better with the transcript abundance on real ribo-seq data.

## 2. Approach

Ribomap works in 3 stages (Figure 1; see also Supplementary Material):

**Figure 1:**
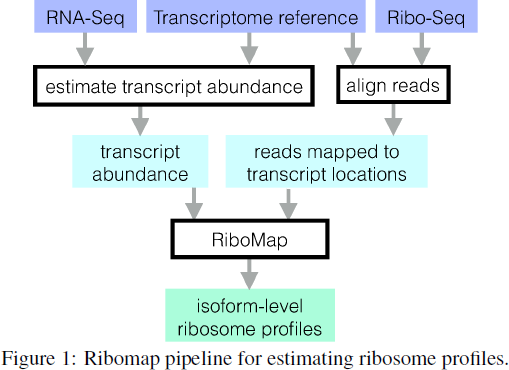
Ribomap pipeline for estimating ribosome profiles.

Transcript abundance estimation. Since RNA-seq experiments should always be performed in parallel with ribo-seq, the abundance *α*_*t*_ per base of each transcript *t* can be estimated from the RNA-seq data using Sailfish [27], an ultra-fast mRNA isoform quantification package. Ribomap also accepts transcript abundance estimations from cufflinks [36] and eXpress [29].
Mapping ribo-seq reads to the reference transcriptome. We obtain all the transcript-location pairs *L*_*r*_ where the read sequence *r* matches the transcript sequence by aligning the entire set of ribo-seq reads *R* to the transcriptome with STAR [6].
Ribosome profile estimation. Let *c*_*r*_ be the number of ribo-seq reads with sequence *r*. Ribomap sets the number of footprints *c*_*rti*_ with sequence *r* that originate from a specific location *i* on transcript *t* to be proportional to the transcript abundance α_*t*_ of transcript 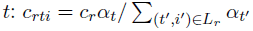, where the denominator is the total transcript abundance with a sequence matching *r*. The total number of reads *cti* that are assigned to transcript *t*, location *i*, is then 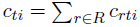. The *c*_*ti*_ give the profiles for each transcript. The sum is needed here because there can exist multiple sequence patterns being mapped to the same transcript location due to sequencing errors, so the final estimated ribosome count for a transcript location should be the sum of the estimated count for all matched sequence patterns.

## 3. Results and Discussion

To evaluate the performance of Ribomap, we generated simulated ribo-seq reads with known ground truth profiles using transcript abundance of GSM546921 RNA-seq data [11] and a dynamic range of initiation rates. Ribosome occupancy probabilities for locations on a given transcript were simulated using the ribosome flow model [28]. Errors were added to the reads using a Poisson process with a rate of 0.5*%*, which was estimated from the ribo-seq data GSM546920 [11]. For comparison, we also test a naive approach, called “*Star prime*”, that maps each read to a single candidate location. More details are in Supplementary material.

The Pearson correlation coefficients between Ribomap’s ribosome profiles and the ground truth is significantly higher than that of Star prime (Figure 2): 81% of our profiles have a higher Pearson correlation (Mann-Whitney U test *p* < 3×10^308^) and 68% have a smaller root mean square error (Mann-Whitney U test *p* = 3.3×10^221^). This suggests that Ribomap more accurately recovers the ribosome profiles than the standard mapping procedure applied to isoforms.

**Figure 2:**
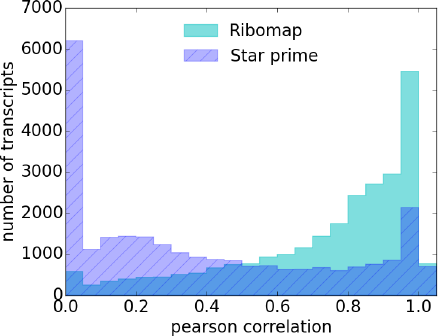
Histogram of the Pearson correlation between the footprint assignments and the ground truth profiles. Ribomap has a significant higher Pearson correlation (median: 0.83) than Star prime (median: 0.28). The spike at 0 of Star prime is due to STAR not assigning footprints to transcripts that are estimated to be present.

The good correlation between the ground truth profile and the estimated profile also leads to a good estimation of the total ribosome loads on a transcript. Ribomap’s ribosome loads estimation on non-simulated ribo-seq data (GSM546920, [11]) correlates well with the estimated transcript abundance (Pearson *r* = 0.71). We do not expect a perfect correlation due to isoform-specific translational regulation. On the other hand, the pure mapping-based approach of Star prime does not correlate as well (*r* = 0.28).

Through two lines of evidence, on real and simulated ribo-seq data, we show that Ribomap produces useful, high-quality ribosome profiles along individual isoforms. It can serve as a useful first step for downstream analysis of translational regulation from ribo-seq data.

## Supplementary Information

## 4. More details on the Ribomap pipeline

### 4.1 Background

Quantifying ribosome occupancy correctly is the first step of analyzing the ribo-seq data. Measurements of protein translation efficiencies [11], ribosome loads [14], pileups and stalling [15] are all derived from ribosome profiles. The challenges of generating such a profile includes deconvolving multi-mapped reads, selecting the correct codon location that the P-site maps to, and bias correction [13]. While none of these issues has a standard protocol, we describe Ribomap, an automatic pipeline that addresses the above challenges and outputs isoform-level ribosome profiles and other ribo-seq analyses.

Current approaches to ribo-seq analysis either discard ambiguously mapped reads [11, 14, 24, 35] or randomly assign them to one of the candidate regions [20, 30]. Such approaches can result in an inaccurate estimation of the ribosome profiles. For example, regions without estimated ribosome footprints are indistinguishable from read-free regions caused by discarded multi-mapping reads. Moreover, randomly assigned reads might cause some regions to have a faulty peak in the profile, while leaving other regions footprint-free (Supplementary Figure 3). Reads are therefore generally mapped to genes by choosing a single isoform [11], or by using the union of all possible exons [5, 25, 34]. Although there is a pipeline built to process ribo-seq reads for identifying protein sequences [3], a method proposed to resolve multi-mapping problems caused by repetitive sequences [4] and software developed to align short reads to splice junctions [22], none of these approaches so far handles multi-mapping problems caused by alternative splicing, which, as is shown in the main text and is further shown here in Supplementary Figure 4, is the major cause of ambiguous mappings. Without dealing with such source of multi-mappings, the current approaches can only be used to estimate the overall ribosome abundance of a given gene, and are incapable of estimating ribosome profile of different mRNA isoforms. These simplistic approaches can also lead to incorrect gene-level estimates — see Supplementary Figure 5.

The better assignment mechanism described here results in a more precise estimation of the per-mRNA ribosome profiles. And a better estimation of the ribosome profiles can also lead to a better estimation of the total ribosome loads. Even for the case of the mouse data from [15], where approximately only 50% of the reads have multi-mappings, the estimated ribosome loads correlate better to the estimated transcript abundance (Pearson r=0.56) than the naive Star prime approach (Pearson r=0.45). We believe a better estimation of the isoform level ribosome profiles can lead to a better understanding of translational regulation.

**Supplementary Figure 3:**
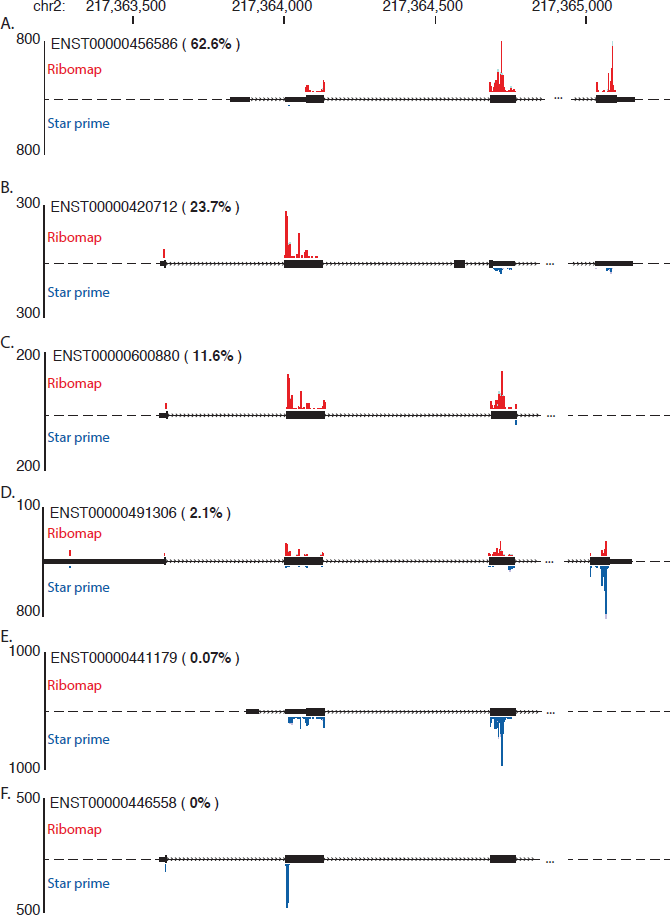
An example of isoform level ribosome profiles for gene RPL37A (ENSG00000197756.5) produced by Ribomap and Star Prime on the Hela cell data (GSM546920) from [11]. The isoforms are listed in order of their relative abundance (value shown next to the transcript ID) estimated from Sailfish [27], and the isoform diagram is shown in between the Ribomap profile and Star prime profile. While Ribomap maps the footprints to the more abundant transcripts, Star prime assigns footprint reads to a single candidate location at random, leaving two of the expressed transcripts almost ribosome-free (A, C), two of the unexpressed transcripts with the highest ribosome loads (E, F), and a surprisingly huge pile-up at the end of transcript ENST00000491306, inconsistent with its surrounding ribosome pile-ups (D). Profile plot is shown in UCSC Genome Browser [17].

**Supplementary Figure 4:**
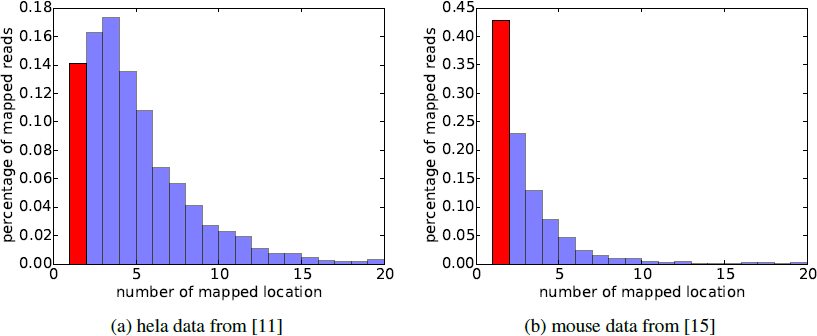
Histogram of the number of mappings for all mapped ribo-seq reads in two data sets. Reads are mapped to the human and mouse transcriptome respectively. The proportion of uniquely mapped reads are marked in red. More than 50% of the reads in both data sets are ambiguous mapped, and ambiguous mapping are extremely common in Human reads. Even for the mouse data, where ambiguous mappings is less prevalent, 68% of the multi-mappings are caused by alternative splicing.

**Supplementary Figure 5:**
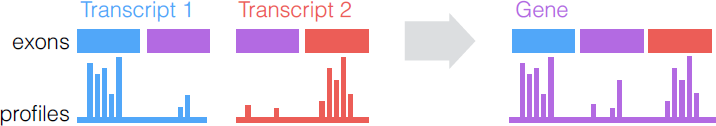
An illustration of when estimating ribosome profile on a gene level will fail. One such case is the pileup does not happen in all isoforms, where merging the isoform-level profile will result in multiple peaks being present simultaneously. Another case is that the pileup in an exon location is not significant in any isoforms, but accumulating them together might produce a faulty peak.

### 4.2 Rationale of the Ribomap footprint assignment scheme

For a ribosome footprint read to come from a transcript location two conditions must hold: first, the transcript has to be present in the cell; second, a ribosome has to be translating the current codon. Therefore, in order to quantify the observed ribosome pileup, we make the following two assumptions: First, identical transcripts have identical translation dynamics; second, each codon location shares a unique pileup behavior due to how fast the current codon can be elongated and the surrounding codon pileups. This means the final observation of the number of ribosome footprints *c*_*mi*_ from transcript *m* at location *i* is proportional to both the chances of observing the specific transcript codon fragment in the cell *α*_*m*_ (transcript abundance per base) and the chances of a ribosome occupying such a codon location *p*_*mi*_ given that the ribosome is from this transcript: *c*_*mi*_ ∝ α*m* × *p*_*mi*_.

**Supplementary Figure 6:**
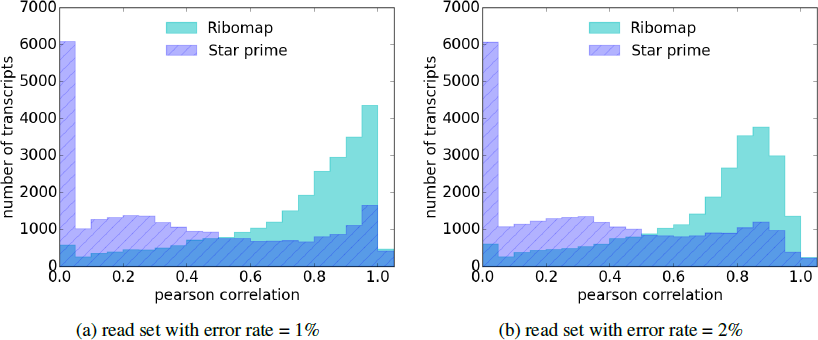
Histogram of the Pearson correlation between the footprint assignments and the ground truth ribosome profiles on read sets with different error rates. Ribomap has a significantly higher Pearson correlation than Star prime (Mann-Whitney-U p< 3×10308 for both cases). For simulated set with error rate 1%, the median Pearson correlation is 0.81 for Ribomap and 0.30 for Star prime; for simulated set with error rate 2%, the median Pearson correlation is 0.78 for Ribomap and 0.31 for Star prime.

The transcript abundance can be estimated from the RNA-seq reads. However, there is no prior knowledge of the per-codon-location specific pileup. We therefore assume the per-codon ribosome abundance is uniformly unbiased. This leads to the assumption that the final ribosome abundance of a transcript location is determined by the transcript abundance: *c*_*mi*_ ∝ *α*_*m*_.

Intuitively, the transcript abundance is like the outline of the profile, and the location-specific ribosome abundance is like the detail of the profile. Our method tries to grasp the outline of the profile first and then let the read sequences themselves take care of the profile details.

### 4.3 Performance under various sequencing error rates

Supplementary Figure 6 gives the analog to Figure 1 in the manuscript, showing the distribution of Pearson correlations under various rates of sequencing error. See Section 6.4 for how these errors were introduced. Under reasonable error rates, the distributions are qualitatively the same.

### 4.4 Additional steps in Ribomap

The raw ribosome profiling reads and the RNA-seq reads are fed into the Ribomap analysis pipeline. Reads are first preprocessed by discarding contaminated reads and trimming the 3’ adapter portion of the reads. The remaining reads are then aligned to the transcriptome to find all possible candidate mapping locations for all reads. The transcript abundances are also estimated from the RNA-seq reads. The last step of ribomap takes in the transcript abundance estimation and the read mappings, and produces a vector of read count per codon position for each transcript. The P-site location for each ribo-seq read are decided dynamically based on the read length. The sequencing bias in the ribosome profiles are corrected by normalizing a transcript’s ribosome profile with its mRNA profile (as done in [1] and [37]). A full description about Ribomap’s output can be found in Section 5. The command to run Ribomap is:

~~~
run_ribomap.sh --rnaseq_fq *rnaseqfq.gz* --riboseq_fq *riboseq.fq.gz* --contaminant_fa *contaminant.fa* --transcript_fa *transcript.fa* --cds_range *cds-range.txt*
~~~

Ribomap uses state-of-the-art read-processing tools for several of its steps. We below list in detail each step of the Ribomap pipeline and the command for executing them, in case the user wants to skip some intermediate steps or make adjustments on individual steps.

#### 4.4.1 Using STAR aligner to filter out contaminating reads

Reads that can be mapped to rRNA, tRNA, snoRNA, may not be representative of ribosome protection *per se* and, in fact, may be contaminants merely associated with the ribosome. As such, they can be filtered out by mapping the reads to the ribosome RNA sequences and transfer RNA sequences of your organism and keeping only unmapped reads for downstream analysis. This is done via the STAR aligner [6].

The first step, which must be done only once per organism, is to create a STAR index for the contaminant RNA sequences with the following command, assuming the sequences of the unwanted molecules are in the file *contamination.fa:*

~~~
STAR --runThreadN *nproc* --runMode genomeGenerate
~~~

~~~
--genomeDir rrna_idx --genomeFastaFiles *contamination.fa*
~~~

~~~
--genomeSAindexNbases 5 --genomeChrBinNbits 11
~~~

*nproc* specifies the number of threads to run STAR. Since STAR treats every sequence entry in the reference FASTA as a ‘genome’, the option `--genomeSAindexNbases 5` forces STAR to build the index properly for small molecule sequences. We include both tRNA and rRNA sequences as contaminants in our analysis.

Once this index is created, it can be used to filter the contaminated reads as follows, assuming the zipped FASTQ file of raw sequencing reads is in *riboseq.fq.gz:*

~~~
STAR --runThreadN *nproc* --genomeDir rrna_idx --readFilesIn *riboseq.fq.gz*
~~~

~~~
--readFilesCommand zcat --outFileNamePrefix *riboaligned*
~~~

~~~
--outStd SAM --outReadsUnmapped Fastx --outSAMmode NoQS
~~~

~~~
--clip3pAdapterSeq *adapter* —seedSearchLmax 10
~~~

--outFilterMultimapScoreRange 0 --outFilterMultimapNmax 255

~~~
--outFilterMismatchNmax *nmismatch*
~~~

~~~
--outFilterIntronMotifs RemoveNoncanonical > /dev/null
~~~

where *adpater* is the adapter sequence (TCGTATGCCGTCTTCTGCTTG for the Hela data set, and `CTGTAGG`

CACCATCAATTCGTATGCCGTCTTCTGCTTGAA for the mouse data set), and *nmismatch* = 1 is the number of allowed mismatches for the alignment. The option `--outReadsUnmapped Fastx` causes STAR to output the unmapped reads to a FASTA file called *riboaligned* `Unmapped.out.mate1`;
`-- seedSearchLmax 10` increases the sensitivity of STAR for aligning short reads;
`--outFilterMultimapScoreRange 0` guarantees that only the alignments with the best scores are reported; `--outFilterMultimapNmax 255` filters out reads that can be mapped to more than 255 locations. STAR by default clips off read ends if a better local alignment score can be achieved. Such a procedure, called ‘soft clipping’, is very useful for handling ribo-seq reads since the first couple of bases in the 5’ end are likely to be contaminated, and an adapter sequence is usually attached to the 3’ end.

#### 4.4.2 Map the remaining reads to the transcriptome

After removing reads that may have been the result of contamination we are now ready to align the remaining reads to the transcriptome. This step is also accomplished by STAR. Assuming the transcriptome sequences are in *transcript.fa,* the command to generate the transcriptome index is:

~~~
STAR --runThreadN *nproc* --runMode genomeGenerate
~~~

~~~
--genomeDir transcript_idx --genomeFastaFiles *transcript.fa*
~~~

~~~
--genomeSAindexNbases 11 --genomeChrBinNbits 12
~~~

Again, `--genomeSAindexNbases 11` insures that the index is built properly for shorter molecules (compared to chromosome), and `--genomeChrBinNbits 12` reduces the memory consumption when there are many reference sequences provided.

The command to align the reads to the transcriptome is:

~~~
STAR --runThreadN *nproc* --genomeDir transcript_idx
~~~

~~~
--readFilesIn *riboaligned*Unmapped.out.mate1
~~~

~~~
--outFileNamePrefix *riboaligned-transcript*
~~~

~~~
--clip3pAdapterSeq *adapter* --seedSearchLmax 10
~~~

~~~
--outFilterMultimapScoreRange 0 --outFilterMultimapNmax 255
~~~

~~~
--outFilterMismatchNmax *nmismatch*
~~~

~~~
--outFilterIntronMotifs RemoveNoncanonical
~~~

~~~
--outSAMtype BAM Unsorted --outSAMmode NoQS
~~~

~~~
--outSAMattributes NH NM
~~~

The aligned reads will be stored in riboaligned_transcript`Aligned.out.bam`. It includes two SAM attributes for each alignment record: `NH` is the number of reported alignments for the read, and `NM` is the number of mismatches in the current alignment.

#### 4.4.3 Transcript abundance calculation using Sailfish [27]

The transcript abundance estimation is used to guide the isoform-level ribosome profile estimation. We are currently using the latest version of Sailfish (called Salmon) that supports the read alignment results as the input. Assuming that the read alignments of the RNA-seq data are in `rnaaligned_transcriptAligned.out.bam`, the command to perform the transcript abundance estimation is:

~~~
salmon quant -t *transcript.fa* -l SF -a
~~~

~~~
rnaaligned_transcriptAligned.out.bam
~~~

~~~
-o sm_quant -p *nproc* —bias_correct
~~~

The flag `--bias_correct` allows Salmon to correct for sequencing biases in the RNA-seq reads. The transcript abundance estimation is in `sm_quant/quant_bias_corrected.sf`.

#### 4.4.4 Isoform-level ribosome profile estimation

This is the last step of the Ribomap analysis pipeline, and it is automatically handled by an executable (developed by us) called `riboprof`. It takes in the transcriptome fasta file, a CDS range file, the ribo-seq and RNA-seq alignment bam files (produced above), the transcript abundance estimation file (produced as above; or it also supports abundance estimations from eXpress [29] or Cufflinks [36] if those estimates are preferred and available). The CDS range file (assume its name is `cds_range.txt`) gives the coding region for each transcript. The command to perform an isoform-level ribosome profile estimation is:

~~~
riboprof --fasta *transcript.fa* --cds_range cds_range.txt
~~~

~~~
--mrnabam rnaaligned_transcriptAligned.out.bam
~~~

~~~
--ribobam riboaligned_transcriptAligned.out.bam
~~~

~~~
--min_fplen *min_fplen* --max_fplen *max_fplen* --offset *offset.txt*
~~~

~~~
--sf sm_quant/quant_bias_corrected.sf —tabd_cutoff *tabd_cutoff*
~~~

~~~
--out *ribomap-out*
~~~

Only reads with size (after ‘soft clipping’) between *min_fplen=25* and *max_jplen=36* are kept for estimating the ribosome profiles. Alignments with RC flag set (reads are reverse complemented and then aligned to the transcriptome) are discarded due to the single strandedness of the ribosome profile protocol. *offset.txt* provides the P-site offset of a read given the read length. Following [15], we assign the P-site of reads with length < 30 to be 12, with length between 31 and 33 to be 13, and with length > 33 to be 14. Only transcripts with abundance greater than *tabd_cutoff* = 0 are considered to be expressed and are included for ribosome profile estimation.

There are three iterations in the footprint assignment procedure: First, only candidate locations with a frame-0 P-site mapping are considered; second, the remaining reads are assigned to frame 1 and 2 locations; third, the rest of the mapped reads - reads that cannot be mapped to any CDS regions, are mapped to UTR regions. For all three iterations, reads are mapped to candidate locations proportional to the transcript abundance if multiple locations are presented for one read.

The estimated ribosome profiles and other analysis will be written to the directory *ribomap_out.*

More information about the options and input file formats for Ribomap can be found in the README file in Ribomap’s Github page (https://github.com/Kingsford-Group/ribomap/blob/master/README.md).

#### 4.5 Settings of Star prime

By default, STAR only marks one alignment for multi-mapping reads as primary (FLAG 0x100 unset), such an alignment “is randomly selected from the alignments of equal quality.” And this primary alignment is used to estimate ribosome profiles in *Star prime.* In addition, no prior transcript abundance knowledge is used in *Star prime*, and therefore all transcripts in the transcriptome are taken into account. All other settings (i.e. contaminated read filtering, adapter clipping, read size selection, dynamic P-site assignment) are kept the same as the Ribomap pipeline described above.

#### 4.6 Running time and memory usage of Ribomap

Ribomap runs for about 15 minutes on the ribo-seq data GSM546920 with 18 million reads on 15 threads. The running time includes the time to build the STAR index for both the contaminated sequences and the transcriptome, filtering the contaminated reads and aligning the remaining reads to the transcriptome for both RNA-seq and ribo-seq data, transcript abundance estimation, and estimating isoform level ribosome profiles. Memory usage is about 8.6G.

## 5. Output file format

In addition to estimated footprint count per codon position for each transcript isoforms, Ribomap also outputs sub-codon resolution, nucleotide-level ribosome profiles including the UTR regions. Furthermore, Ribomap reports ribosome loads, translation efficiency, and the relative abundance for each transcript.

Lastly, Ribomap reports transcripts in order of the rank difference between the relative transcript abundance and ribosome load, to help identify isoforms with different translation efficiency.

Ribomap outputs five files:

File 1: 

~~~
XXX.base
~~~

 gives the sub-codon resolution, nucleotide-level ribosome profiles including the UTR regions. Only transcripts with a non-zero total ribosome count are reported. Each entry of a specific transcript looks like this:

~~~
refID: 0
~~~

~~~
tid: YAL001C
~~~

~~~
ribo profile: 0 0 0 74 68 …
~~~

~~~
mRNA profile: 31 35 50 73 87 96 104 …
~~~

~~~
normalized ribo profile: 0 0 0 1.0137 0.781609 0.0208333 0.125 …
~~~

where:

**refID** is the transcript fai index in *transcript.fa.*

**tid** is the transcript header name in *transcript.fa*.

**ribo profile** nucleotide level ribosome profile including the UTR regions.

**mRNA profile** RNA-seq read coverage profile.

**normalized ribo profile** is the ribosome profile after bias correction. Each number in the vector is the ratio between the ribo profile count and the mRNA profile count.

File 2: 

~~~
XXX.codon
~~~

 gives the in-frame ribosome profiles for each transcript within the CDS region. The file format is the same as the `XXX.base` file.

File 3: 

~~~
XXX.stats
~~~

 is the summarized statistics for each transcript. Each entry of a specific transcript looks like this:

~~~
refID: 0
~~~

~~~
tid: YAL001C
~~~

~~~
rabd: 3959
~~~

~~~
tabd:0.00020934
~~~

~~~
te: 1.89078e+07
~~~

where:

**rabd** is the total ribosome loads, which is the sum of the `ribo profile` vector in `XXX.base`.

**tabd** is the relative transcript abundance from Sailfish’s result.

**te** is the relative translational efficiency, which is the ratio between `rabd` and `tabd`.

File 4: 

~~~
XXX_abundant.list
~~~

 gives a list of transcripts whose total ribosome abundance is more than expected given the transcript abundance. Such a list explores the difference between the ribosome abundance ranking and the transcript abundance ranking. A higher rank of total ribosome footprint count compared to the transcript abundance might indicate that the transcript is more highly packed with ribosomes and this might suggest that there is a translational amplification regulation for this transcript. Each line in the file looks like this:

~~~
ENST00000340756.2 2.81302e-09 1046.59 0 96 -96
~~~

It is the transcript header name followed by several statistics, in order:

1. relative transcript abundance,
2. total ribosome footprint count,
3. the percentile ranking of the transcript abundance,
4. the percentile ranking of the total ribosome loads (transcripts with zero total ribosome loads are excluded from the analysis),
5. difference between the transcript abundance rank and the total ribosome footprint count rank.

The list of entries in this file is ordered by the ranking difference between the transcript abundance and the ribosome abundance. Only transcripts with ranking differences greater than 10 are listed.

File 5: 

~~~
XXX_scarce.list
~~~

 gives alist of transcripts whose total ribosome abundance is less than expected given the transcript abundance. Analogous to the abundant list, this list include transcripts that might have a translational buffering regulation. The file format is the same as 

~~~
XXX_abundant.list
~~~

.

## 6. Generating simulated footprints

To evaluate the performance of the ribosome footprint read assignment, we generated simulated footprint data with known ribosome profiles. Let *N* be the total number of transcripts, *α*_*m*_ be the per-base relative abundance of a transcript *m*, and *p*_*mi*_ be the ribosome occupancy probability of a location *i* on transcript *m*. The number of simulated footprints from location *i* on transcript *m* is set to be *N* ×*α*_*m*_×*p*_*mi*_, where N is set so that a total of 20,000,000 footprints from all locations of all transcripts are generated. How *α*_*m*_ and *p*_*mi*_ are set is described below.

### 6.1 Transcript abundance

The transcript abundance is estimated from the RNA-seq data paired with the ribosome profiling experiments (GSM546921) via Sailfish [27]. Only transcripts with TPM value (transcripts per million) greater than 1 are included, which results in a total of 39,414 transcripts. The relative transcript abundance per base *α*_*m*_ is computed as follows: Let *t*_*m*_ be the transcript abundance estimated from Sailfish for transcript *m*, let *l*_*m*_ be the transcript length, let *T* be the transcriptome, 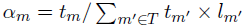.

The rationale of using transcript abundance per base for assigning the reads is that each transcript codon location can be seen as a type of ball with a unique color, and the probability of observing such a transcript codon fragment in the cell can be compared as randomly picking a ball of a specific color from a pool of balls. We assume codon locations on the same transcript have equal visibilities. For the extreme case where there is only one type of transcript in the cell with *l*_*m*_ bases, the probability of seeing any codons is 1/*l*_*m*_. For the case where there are more than one type of transcript, if there are *t*_*m*_ transcripts with type *m*, then there will be *t*_*m*_ copies of each codon positions, and the probability of seeing a specific codon position on transcript *m* over all possible codon positions is: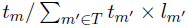.

### 6.2 Ribosome occupancy probability

We model the movement of ribosomes on a given transcript with a ribosome flow model based on a total asymmetric exclusion process (TASEP) [28]. In this process, each mRNA is modeled as a sequence of codons, with ribosomes moving along it with codon-specific elongation rates. The model is “asymmetric” because the ribosome can only move from the 5′ end to the 3′ end of the mRNA. The model has “exclusion” because a location can only be occupied by one ribosome at a time; a ribosome can only move from location *i* to the next one when it is currently at location *i* and the next location is not occupied by another ribosome.

For a mRNA m of length n, the entire translation process is modeled in three steps: First, the ribosome binds to the the start codon of the mRNA with initiation rate *λ*_*m0*_. Second, the ribosome moves along the mRNA from codon *c*_*i*_ to *c*_*i*+1_ with a elongation rate *λ*_*mi*_ (*i* = 1… *n*−1). Third, the ribosome terminates when it reaches the the stop codon *c*_*n*_ with a termination rate *λ*_*mn*_, and the whole peptide chain of the protein will be formed. The *λ* vector thus describes the transition rates of the ribosome moving from one location to another.

The model states, when given a transcript *m*, that the change of the probability of observing a ribosome on location *i* (*p*_*mi*_) is the difference between the incoming and the outgoing flow of the ribosome, modeled using the differential equations:

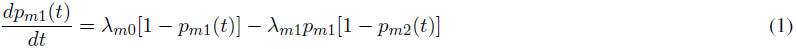

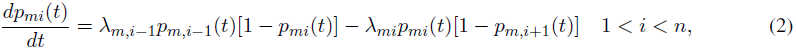

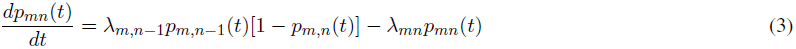

Equation (1) and (3) describe the boundary case of the process, and equation (2) describes the intermediate case.

The ribosome profile under this model is the ribosome occupancy probability distribution when the steady state of the model is reached, during which the probability of observing a ribosome at any location will not change over time. This stationary distribution probability can be solved by setting the left hand side of the equations above to zero.

To generate simulated profiles, we solve the above equations to find the steady state for all 39,414 transcripts. This profiles the probabilities *p*_*mi*_ that a read is drawn from location *i* on one copy of transcript *m*. Selection of the elongation rates (*λ*_*mi*_) and initiation rates (*λ*_*m*0_) is described below.

#### 6.2.1 Elongation rate

Following Reuveni *et al.* [28], we assume the elongation rate is codon-specific and is proportional to the tRNA abundance of a given codon. We use tRNA gene copy number as an approximation to the tRNA abundance in the cell, and we set the elongation rate to be the absolute adaptiveness value (*W*_*i*_) of a given codon *i*:

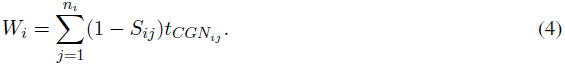

Here *n*_*i*_ is the number of tRNA isoacceptors recognizing codon *i*. *tCGN*_*ij*_ is the gene copy number of the *j*th tRNA that recognizes the ith codon, and *S*_*ij*_ is the selective constraint on the efficiency of the codon-anticodon coupling by considering the wobble paring [7]. The absolute adaptiveness of a given codon gives an estimate for the total amount of tRNAs that can match to a specific codon. The larger the adaptiveness value is, the more efficient a codon can be translated, and thus the faster the elongation rate will be. The elongation rates range between 3.5 and 33 time steps from this calculation.

#### 6.2.2 Initiation rate

The initiation rate is thought to be the rate limiting step during the translation process [14]. We set the initiation rate range to be approximately 100 times smaller than the elongation rate as in [31], and the rates are uniformly sampled for each transcript between 0.03 and 0.3 timesteps.

### 6.3 Ribosome footprint generation

We calculate *α*_*m*_ and *p*_*mi*_ for every transcript as described above and then sample reads by selecting a random transcript with probability proportional to its *α*_*m*_ and then selecting ribosome position *i* proportional to *p*_*mi*_. We then extract a 30-bp read with the 12^*th*^ position set to the ribosome P-site.

### 6.4 Introducing sequencing errors in the read set

We estimate the frequency of sequencing errors from ribosome profiling data (GSM546920 [11]) to be the total number of mismatches of read assignments in the data over the total number of aligned bases (total number of aligned reads × read length). The error rate estimated on the ribosome profiling data GSM546920 is 0.5%.

To add simulated sequencing errors to our simulated reads, we apply a Poisson process to choose the bases to mutate in the simulated footprint read set with the rate parameter set to the error rate. We also tried different error rates (0.5%, 1%, 2%), which all produce similar results in the footprint assignments (Supplementary Figure 6). The histogram in the paper is generated from the data with error rate 0.5%.

### 6.5 RNA-seq data generation

We generated simulated RNA-seq reads with the following procedure: The transcript abundance estimation from Section 6.1 is used as a prior to generate 20,000,000 simulated RNA-seq fragments with default parameters using rlsim [32]. Consistent with the ribosome footprint generation settings, only transcripts with TPM value (transcripts per million) greater than 1 are included. We then truncate the fragments into 36bp-long single-end reads.

When applying Ribomap, we re-estimate the transcript abundance as usual from the simulated RNA-seq reads via Sailfish [27] without looking at the original estimates from the true data.

The analysis shown in Figure 2 in the main text and in Supplementary Figure 6 only uses transcripts with non-zero total ribosome counts, which results in 26,297 transcripts included.

## 7. Data used in the experiments reported here

- **Hela ribosome footprint data:** Human Hela cell ribo-seq mock 32hr runs1-2 in [11] (GSM546920).
- **Hela RNA-seq data:** Human Hela cell RNA-seq mock 32hr runs1-3 in [11] (GSM546921).
- **Human Transcriptome reference fasta file:** The human protein-coding transcriptome fasta file is downloaded from the Gencode website: (ftp://ftp.sanger.ac.uk/pub/gencode/Gencode_human/release_18/gencode.v18.pc_transcripts.fa.gz). The CDS region information used in our analysis is obtained from the headers of the fasta sequence entries.
- **Human Transcriptome gene annotation gtf file:** The human gene annotation gtf is also downloaded from the Gencode website: (ftp://ftp.sanger.ac.uk/pub/gencode/Gencode_human/release_18/gencode.v18.annotation.gtf.gz). This file is used to obtain the frame information of the CDS regions of the transcripts.
- **Mouse Ribosome footprint data:** ES cell feeder-free, w/ LIF 60 s CYH (100 ug/ml) ribo_mesc_yeslif Illumina GAIIin [15] (GSM765301).
- **Mouse RNA-seq data:** ES cell feeder-free, w/ LIF 60 s CYH (100 ug/ml) mrna_mesc_yeslif Illumina GAII in [15] (GSM765289).
- **Mouse Transcriptome reference fasta file:** The mouse protein-coding transcriptome fasta file is downloaded from the Gencode website: (ftp://ftp.sanger.ac.uk/pub/gencode/Gencode_mouse/release_M4/gencode.vM4.pc_transcripts.fa.gz). The CDS region information used in our analysis is obtained from the headers of the fasta sequence entries.
- **Mouse Transcriptome gene annotation gtf file:** The mouse gene annotation gtf is also downloaded from the Gencode website: (ftp://ftp.sanger.ac.uk/pub/gencode/Gencode_mouse/release_M4/gencode.vM4.annotation.gtf.gz). This file is used to obtain the frame information of the CDS regions of the transcripts.
- **Human tRNA gene copy number:** Downloaded from the gtrna database [2]: (http://gtrnadb.ucsc.edu/Hsapi/Hsapi-summary-codon.html).
- **Contaminated sequences:** Both ribosomal sequences and tRNA sequences are included in the contaminated sequences. Ribosomal sequences are downloaded from Ensemble’s [8] ncRNA database with gene_biotype as rRNA; tRNA sequences are downloaded from the gtrna database [2] (http://gtrnadb.ucsc.edu/download/tRNAs/eukaryotic-tRNAs.fa.gz).

